# Flagellin in the human gut microbiome is a diet-adjustable adjuvant for vaccination

**DOI:** 10.1101/2025.02.21.639485

**Authors:** Kelsey E. Huus, µHEAT Clinical Study Group, Yi Han Tan, Hirohito Abo, Ronald Keller, Ezgi Atay, Dai Long Vu, Silke Dauser, Alina Prokipchuk, Alexander V. Tyakht, Nicholas Youngblut, Benoit Chassaing, Sang-Moo Kang, Julie Parsonnet, Peter G. Kremsner, Lisa Maier, Andrew T. Gewirtz, Meral Esen, Ruth E. Ley

**Affiliations:** Department of Microbiome Science, Max Planck Institute for Biology, Tübingen, Germany; Controlling Microbes to Fight Infections (CMFI) Cluster of Excellence, Tübingen, Germany; Institute for Tropical Medicine, University of Tübingen, Tübingen, Germany; Institute for Biomedical Sciences, Georgia State University, Atlanta, USA; Department of Microbiology, Pasteur Institute, Paris, France; Department of Medicine, Stanford University, Palo Alto, USA; Centre de Recherches Medicales de Lambaréné, Lambaréné, Gabon; Interfaculty Institute of Microbiology and Infection Medicine, University of Tübingen, Tübingen, Germany

## Abstract

The intestinal microbiota is thought to modulate immune responsiveness to vaccines. Human studies on this topic, however, have yielded inconsistent results^1,2^. We hypothesized that the microbiome would influence innate immune responses, and thus vaccine reactogenicity, more directly than vaccine immunogenicity. To test this, we established the µHEAT (Microbial-Human Ecology And Temperature) study, which longitudinally profiled the fecal microbiota, oral body temperature and serum antibody responses of 171 healthy adults (18-40 years old) before and after vaccination for SARS-CoV-2. Increased temperature after vaccination (ΔT) was associated with habitual diet and with baseline metabolic and immune markers. The microbiomes of ΔT-high (ΔT^hi^) participants were characterized by high expression of flagellin and an overabundance of the flagellated bacterium *Waltera*. Fecal samples from ΔT^hi^ participants induced more inflammation in human cells and stronger post-vaccine temperature responses in mice compared to ΔT^lo^ samples, suggesting a causal role for the microbiome. Moreover, *Waltera* flagellin replicated the inflammatory phenotypes *in vitro* and was modulable via a dietary additive. Overall, these data suggest that flagellin from the gut microbiome stimulates innate immunity and vaccine reactogenicity, and that this axis can be manipulated via diet. These findings have implications for improving human vaccine tolerance and immunogenicity.

## INTRODUCTION

Vaccination is a crucial public health tool for disease prevention^3^. However, vaccine efficacy varies widely between people and across populations^4,5^. While multiple factors contribute to variation in vaccine responses, including host genetics, age and sex, the intestinal microbiome has gained significant interest for its potential as a targetable modulator of vaccine responses^1,2^. The gut microbiome was found to influence antibody responses to flu vaccine in mice and humans^6–8^, and variations in human microbiome composition have been associated with antibody titers to multiple human vaccines, including SARS-CoV-2^9–13^. Despite these efforts, no consistent microbiome signature of vaccine responsiveness has emerged across studies, and the importance of the gut microbiome in human vaccine immunogenicity remains debated.

One mechanism by which the microbiome might influence vaccine responses is the ‘natural adjuvant’ hypothesis, whereby the microbiome stimulates innate immunity, thus promoting vaccine-specific adaptive immune responses^1^. For example, microbiome flagellin signalling through toll-like receptor 5 (TLR5) promoted antibody responses to non-adjuvated vaccines in mice^6^. These data suggest that microbiome control of vaccination acts at the level of innate immunity. However, nearly all human microbiome-vaccine studies have focused on antibodies as the primary outcome. Existing studies may therefore have missed important microbiome-vaccine associations when antibody levels are influenced by other factors downstream of innate immunity (for example, by prior exposure and memory responses).

We hypothesized that the microbiome influences innate immune responses to vaccination, and that this can best be measured by studying vaccine reactogenicity (vaccine adverse reactions) rather than immunogenicity (antibody titres). Mass vaccination campaigns against SARS-CoV-2 provided an ideal model system, as these vaccines commonly induced systemic side effects including fever, and fever represents a broad, accessible and easily quantifiable read-out of innate immunity. To this end, we established the µHEAT (<u>Micro</u>bial-<u>H</u>uman <u>E</u>cology and <u>T</u>emperature) study, which longitudinally profiled oral body temperature, serum immune markers and fecal microbiome in healthy adults before and after SARS-CoV-2 vaccination (Fig 1A). We find that production of flagellin within the human gut microbiome associates with increased body temperature after vaccination, drives host inflammation, and may be modulated by diet.

**Fig 1.**
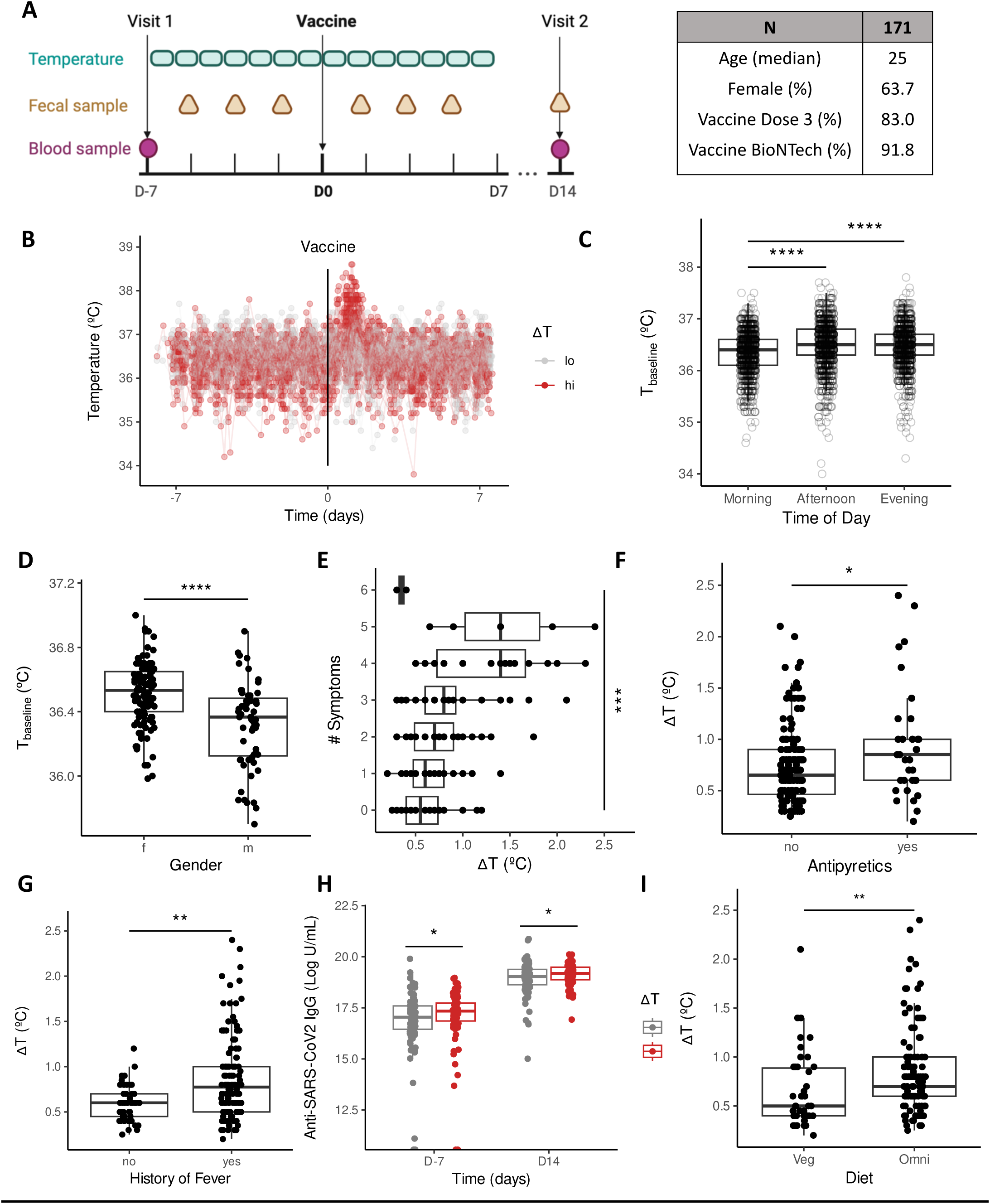
Vaccine-induced thermogenesis is associated with habitual diet. (A) Schematic of the µHEAT study protocol and basic participant demographics. (B) Oral body temperature recorded three times per day per participant (N=7079 records from 171 people). (C) Baseline (pre-vaccine) temperature records according to time of day at measurement. (D) Median baseline temperature per person by gender. (E-G) Degree of post-vaccine temperature increase (ΔT) versus (E) total number of symptoms reported by the participant post-vaccine, (F) reported consumption of antipyretics post-vaccine and (G) history of fever, i.e., whether the participant reported experiencing any fever to prior vaccines or illnesses in the past two years. (H) Serum anti-SARS-CoV-2 IgG pre– and post-vaccine, according to ΔT category. (I) ΔT according to self-reported habitual diet. Statistical significance was determined via ANOVA with post-hoc Tukey’s test (C), Wilcoxon Rank-Sum test (D, F-I) or Kruskal-Wallis test (E).

## RESULTS

### Vaccine-induced thermogenesis is associated with habitual diet

The µHEAT study followed 171 individuals before and after mRNA vaccination for SARS-CoV-2, the majority of whom received third-dose booster vaccinations (Fig 1A, Tables S1-S2). Participants provided three body temperature records daily over a two-week period (total N=7079 temperature records; Fig 1B). As expected^14^, baseline body temperature showed a strong circadian rhythm (Fig 1C) and was higher on average in women than in men (Fig 1D). Moreover, there was a pronounced spike in participant body temperature following vaccination (Fig 1B).

We defined ΔT as the increase in body temperature post-vaccine, relative to each participant’s median temperature pre-vaccine for that time of day (see Methods). For categorical analysis, we divided participants into ΔT^hi^ (N=72) and ΔT^lo^ (N=99) categories based on the group mean; functionally, this resulted in participants being classified as ΔT^hi^ if their temperature rose 0.8°C or more above personal baseline. Unlike baseline body temperature, ΔT was unaffected by gender or time of day (Fig S1A-B). Further, ΔT did not differ significantly by type or dose of vaccine received during the study, nor by prior vaccine or infection history (Table S3).

ΔT correlated positively with the number of flu-like symptoms reported by the participant after vaccination (Fig 1E) and with subsequent consumption of antipyretic medication (Fig 1F). Participants who reported a fever to any prior dose of the SARS-CoV2 vaccine also measured higher ΔT during the µHEAT study (Fig 1G). Furthermore, ΔT^hi^ participants had higher anti-SARS-CoV-2 antibody titres both before and after booster vaccination (Fig 1H). Together, these data support the use of ΔT to reflect vaccine responsiveness and, consistent with prior reports, indicate that some individuals are consistently prone to stronger fever and antibody responses following SARS-CoV-2 vaccination^15–23^.

Remarkably, we observed that participants who habitually consumed a plant-based diet (self-reported vegans and vegetarians, N=46) had a significantly lower ΔT compared to omnivores (Fig 1I). ΔT trended lower in vegans compared to vegetarians, as well as in vegetarians compared to pescatarians and omnivores (Fig S1C). In contrast, antibody levels were similar by diet both before and after vaccination, suggesting that diet-associated ΔT did not necessarily translate into altered adaptive immunity (Fig S1D). The diet-ΔT association remained significant in linear models controlling for gender, age, BMI, vaccine dose and type, antipyretic usage, and prior immune exposures via both prior fever and starting antibody levels (p=0.004, Table S3). In summary, those who followed a plant-based diet experienced less vaccine-induced thermogenesis.

### **ΔT^hi^** individuals have altered metabolic and immune markers

To better understand the molecular signatures that underlie a higher risk of vaccine-induced thermogenesis, we performed aptamer-based profiling of 1500 human blood proteins in baseline participant serum (Table S4). This yielded associations with both metabolic and immune proteins (Fig 2A). The protein most significantly increased in ΔT^hi^ participants was apolipoprotein E (apoE), a major cholesterol transporter, and this hit remained significant when adjusting for reported diet (Table S5). We therefore extended our profiling to blood lipids via untargeted mass-spectrometry based metabolomics and lipidomics (Fig 2B, Table S6). This revealed a number of circulating lipids that were more abundant in ΔT^hi^ participants at baseline, including two compounds annotated as ethyl linoleate, a precursor of arachidonic acid which is used in the biosynthesis of inflammatory prostaglandins^24^. Furthermore, targeted quantification of cholesterol levels revealed that the ratio of LDL to HDL cholesterol was increased in ΔT^hi^ participants, indicating a less healthy cholesterol profile at baseline (Fig 2C). These data indicate that there is a systemic metabolic signature associated with the risk of vaccine-induced thermogenesis, consistent with the observed dietary trends^25^.

**Fig 2.**
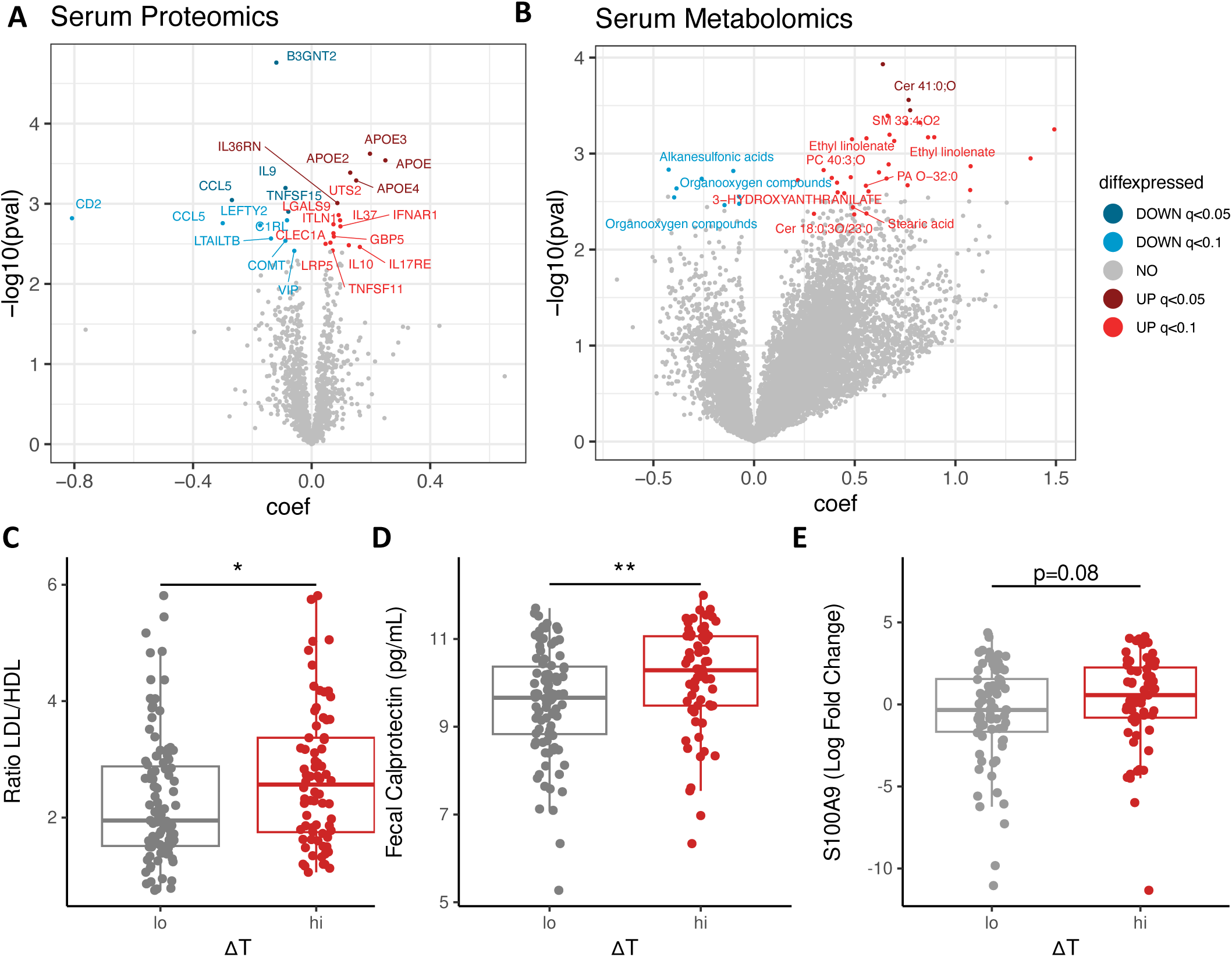
ΔT^hi^ individuals have altered metabolic and immune markers. (A-B) Volcano plot of serum proteins (A) and metabolites (B) significantly enriched (positive coefficient) or depleted (negative coefficient) in ΔT^hi^ participants compared to ΔT^lo^ participants at day D-7. (C) Ratio of LDL to HDL cholesterol in baseline serum of ΔT^hi^ and ^lo^ participants. (D) Fecal calprotectin at day D14 in ΔT^hi^ and ^lo^ participants. (E) Expression of the S100A9 calprotectin subunit in stool at D-1 prior to vaccination. Statistical significance was determined via MaAsLin2 linear models, taking age and sex as covariates, and adjusting p-values via FDR (A-B); or via Wilcoxon Rank-Sum test (C-E).

Contrary to expectations, there was also an overall anti-inflammatory skew in ΔT^hi^ participants, with higher levels of anti-inflammatory cytokines such as IL-10, IL-37 and IL-36RN, and lower levels of pro-inflammatory markers such as the chemokine CCL5, in baseline serum samples (Fig 2A). This immune signature might reflect an attempt by the host to control prior or ongoing inflammation. Furthermore, several proteins with possible roles in microbial control, including IL-17RE (important for intestinal barrier integrity)^26^, and ITLN1 (intelectin-1, an intestinal lectin which binds bacterial glycans)^27^, were significantly increased at baseline in ΔT^hi^ participants.

We thus further tested whether intestinal inflammation was also altered by ΔT category. Strikingly, ΔT^hi^ participants had significantly higher levels of fecal calprotectin two weeks after vaccination (Fig 2D), indicating that ΔT-associated inflammation is also experienced locally within the gastrointestinal tract. Since the preservation of pre-vaccination stool samples in an RNA buffer did not allow for direct calprotectin measurements, we also measured transcripts of the calprotectin subunit S100A9 via qPCR to determine whether intestinal inflammation preceded vaccination. These data suggest that fecal calprotectin is also elevated in ΔT^hi^ participants prior to vaccination (Fig 2E).

Taken together, these data reveal a baseline immunometabolic signature associated with the risk of vaccine-induced thermogenesis. Importantly, ΔT^hi^ individuals also show elevated levels of intestinal inflammation, implicating a possible role for innate immune signalling via the gut microbiome.

### Temperature and vaccination do not impact the intestinal microbiome

To determine whether the gut microbiome associates with vaccine-induced thermogenesis, we generated shotgun metagenomic sequencing from six longitudinal fecal samples per participant over the two-week period before and after vaccination (N=1022 total samples, Fig 3A). We also generated longitudinal fecal transcriptomics from a subset of N=38 participants to assess the functional activity of the microbiome (N=230 samples, Fig 3B; see Methods for subset selection). As expected, people had individualized and relatively stable microbiomes, with longitudinal samples from the same individual more similar to one another than to samples from unrelated individuals, at both the overall community level (Fig 3C) and the level of individual strains (Fig S2 A-B).

**Fig 3.**
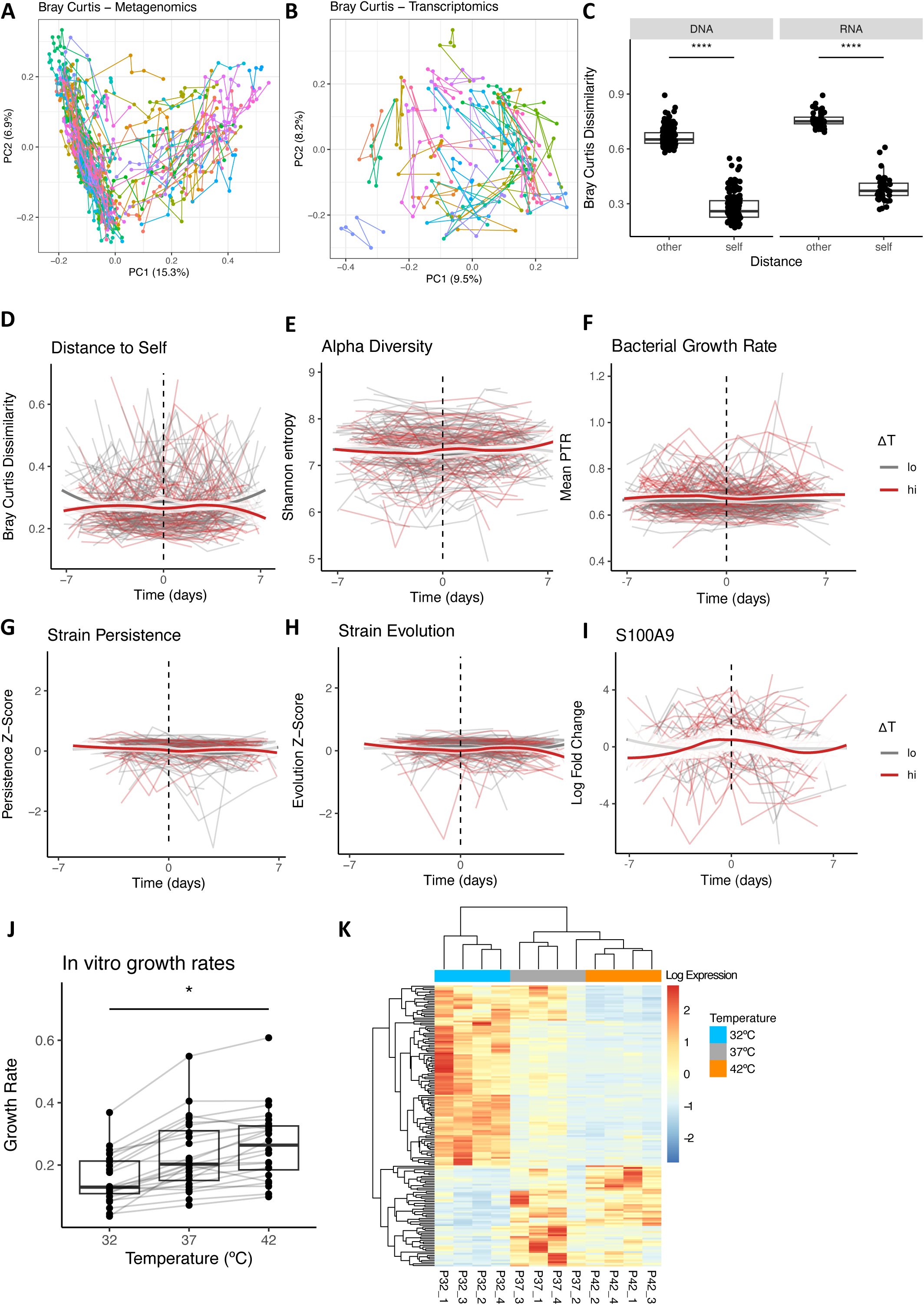
Temperature and vaccination do not impact the intestinal microbiome. (A-B) PCoA plot by Bray-Curtis distance of taxonomic profiles in (A) fecal metagenomic samples (N=942 samples from 171 people after rarefying to 5M reads) and (B) fecal transcriptomic samples (N=226 samples from 38 people after rarefying to 5M reads). Lines join samples per subject. (C) Bray-Curtis dissimilarity quantified between samples of the same individual over time (self) and samples belonging to unrelated individuals (other), for metagenomic (DNA) and transcriptomic (RNA) data. (D) Bray-Curtis dissimilarity of each sample compared to every other sample from that individual, over time. (E-H) Temporal dynamics in metagenomic samples of (E) alpha diversity, (F) average estimated bacterial growth rate (peak-to-trough ratio, PTR), (G) strain persistence, defined as the popANI-based strain similarity in sample X compared to sample 1 and (H) strain evolution, defined as the popANI-based strain similarity in sample X compared to sample X-1. (I) Detection of S100A9 via qPCR in a subset of fecal samples over time (N=298 samples from 47 participants). (J) *in vitro* bacterial growth rates estimated per species, per temperature for a panel of prevalent human gut commensals (N=25). (K) RNA-seq heatmap for genes significantly different after 1 hour of temperature shock in *Segatella copri.* Statistical significance was determined via Wilcoxon Rank-Sum test (C), Kruskal-Wallis test (J) and DESeq2 models with a fold-change cutoff of 1.5 and a FDR-corrected p-value-cutoff of 0.05 (K). Temporal trends in D-I are non-significant (p>0.05) according to mixed linear models using sequencing depth, time, ΔT, and time:ΔT interaction as fixed effects and participant ID as a random effect, with the exception of a slight decrease in strain persistence over time independent of ΔT (p=0.03, see Table S3).

We first tested whether increased body temperature might itself affect the microbiome, hypothesizing that fever-like temperatures or inflammation would be stressful to commensal bacteria. However, we did not observe any changes in microbiome alpha or beta diversity following vaccination, regardless of ΔT category (Fig 3D-E, Fig S2C-E, Table S3). Predicted bacterial growth rates were also stable (Fig 3F), as were measures of strain-level persistence and evolution, with the exception of a slight overall decrease in strain persistence over time (Fig 3 G-H). There was also limited evidence of any individual genera or gene-level transcription changing in abundance in ΔT^hi^ individuals after vaccination (Fig S2F-G, Table S7).

This was initially surprising to us, as fever is commonly presumed to limit microbial growth^28,29^. However, longitudinal assessment of fecal calprotectin levels via qPCR indicated that there was no increase in intestinal inflammation associated with vaccination (Fig 3I). In addition, *in vitro* growth assays of 25 prevalent human gut bacteria indicated that while growth is significantly limited at 32°C compared to 37°C, it increases on average up to 42°C (a physiologically unlikely extreme which exceeds the 38.6°C maximum in our study) (Fig 3J). Even the most temperature-sensitive bacterium in our screen, *Segatella copri,* showed only limited transcriptional changes *in vitro* when shifted from 37°C to 42°C, compared to a more robust response to below-host temperatures (Fig 3K). We therefore concluded that fever-like temperatures do not meaningfully impact gut commensals *in vitro,* and that short-term increases in body temperature are not associated with changes in intestinal inflammation or in the human gut microbiome *in vivo*.

### ΔT^hi^ microbiome is characterized by high flagellin expression and abundance of *Waltera* spp

Having observed that elevated temperatures do not impact the microbiome, we next asked whether the reverse might be true. We proceeded with an analysis of the gut microbiome by ΔT category, using all longitudinal samples per person for greater statistical power, and leveraging mixed linear models via MaAsLin2^30^.

Strikingly, transcriptomic data indicated that multiple flagellins and motility-associated proteins were upregulated in ΔT^hi^ participants across all time points (Fig S3A, Fig 4A, Table S8). This signature was evident when considering RNA relative abundance (Fig S3A-B) but became even more pronounced when RNA abundances were normalized to the respective DNA abundances for that gene and sample, indicating a real transcriptional upregulation (Fig 4A, Fig S3B). Elevated flagellin expression in ΔT^hi^ compared to ΔT^lo^ participants occurred in pre-vaccination samples and remained stable over time (Fig 4B).

**Fig 4.**
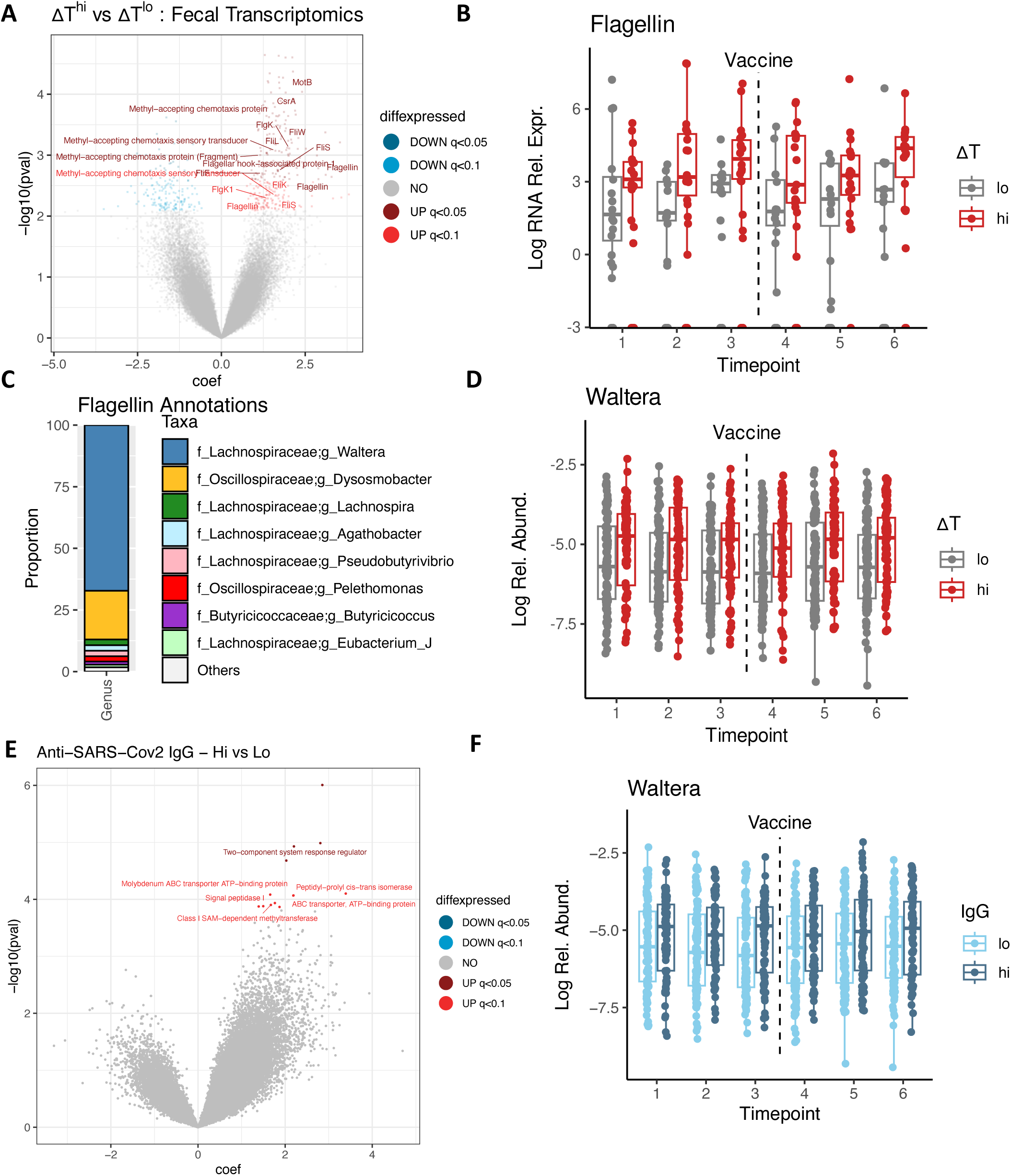
The ΔT^hi^ microbiome is characterized by high flagellin expression and abundance of *Waltera* spp. (A) Volcano plot of genes (UniRef50) significantly enriched and depleted at the transcriptional level, relative to DNA abundance for that same gene, in ΔT^hi^ compared to ΔT^lo^ individuals. Genes involved in motility are labelled. (B) Temporal dynamics of the expression of Flagellin with UniRef50 ID R6ZZ21, one of the most highly differentially expressed flagellins, by ΔT (q<0.05). Log RNA Rel. Expr. is the log-normalized ratio of RNA abundance relative to DNA abundance for that gene. (C) Taxonomic annotations for all flagellins that were significantly differentially expressed by ΔT according to taxa-stratified analysis. (D) Relative abundance of the genus *Waltera* at the metagenomic level, by ΔT over time (q<0.05). (E) Volcano plot of genes significantly enriched and depleted at the transcriptional level, relative to DNA abundance for that same gene, in IgG-high compared to IgG-low individuals (defined by anti-SARS-CoV-2 IgG titres at D14 that are above or below the group mean, respectively). (F) Relative abundance of the genus *Waltera* at the metagenomic level, by antibody category over time (p=0.058). Statistical significance was determined via MaAsLin2 linear models taking sequencing depth, Bristol Stool Score, and time at room temperature as technical covariates, treating participant ID as a random effect, and adjusting p-values via FDR (A-F).

A taxon-stratified analysis revealed that a large portion (>50%) of the upregulated flagellins were expressed by the genus *Waltera*, a *Lachnospiraceae* member, with the remainder assigned to other members of the Bacillota (formerly Firmicutes) phylum (Fig 4C, Table S8). *Waltera* was also more abundant in ΔT^hi^ participants according to metagenomic data, and this signature was also stable over time (q=0.02, Fig 4D, Table S9). Thus, the microbiomes of ΔT^hi^ participants are characterized by a high relative abundance of flagellated *Waltera* and by increased transcriptional expression of commensal flagellins.

In contrast, very few genes were changed at a transcriptional level in high-versus low-antibody participants, and none of these genes were related to flagellins or motility (Fig 4E, Table S10). Although *Waltera* abundance also trended higher in high-antibody individuals over time, this association did not reach statistical significance (Fig 4F, raw p=0.058 by mixed linear model), nor was *Waltera* correlated with the fold-change increases in antibodies with vaccination (p=0.6). Therefore, although higher ΔT is associated with higher antibody responses to vaccination (Fig 1F), the *Waltera-*flagellin signature is more directly associated with ΔT, and would have been missed by a more traditional antibody-focused analysis.

In summary, *Waltera* is overabundant in ΔT^hi^ individuals pre and post vaccination and expresses high levels of transcripts related to flagellar motility.

### The ΔT^hi^ microbiome drives flagellin-dependent inflammation

Bacterial flagellin is a canonical stimulator of innate immunity, and might therefore contribute to the increased inflammation observed in ΔT^hi^ participants. Consistent with this hypothesis, human colonic organoids stimulated with ΔT^hi^ fecal supernatants produced significantly more pro-inflammatory IL-8 compared to stimulation with ΔT^lo^ fecal samples (Fig 5A). IL-8 is produced downstream of TLR5 activation^31^; consistently, organoids also upregulated genes related to TLR signalling cascades (Fig 5B, Table S11). This indicates that fecal bacterial products from ΔT^hi^ individuals can drive inflammation.

**Fig 5.**
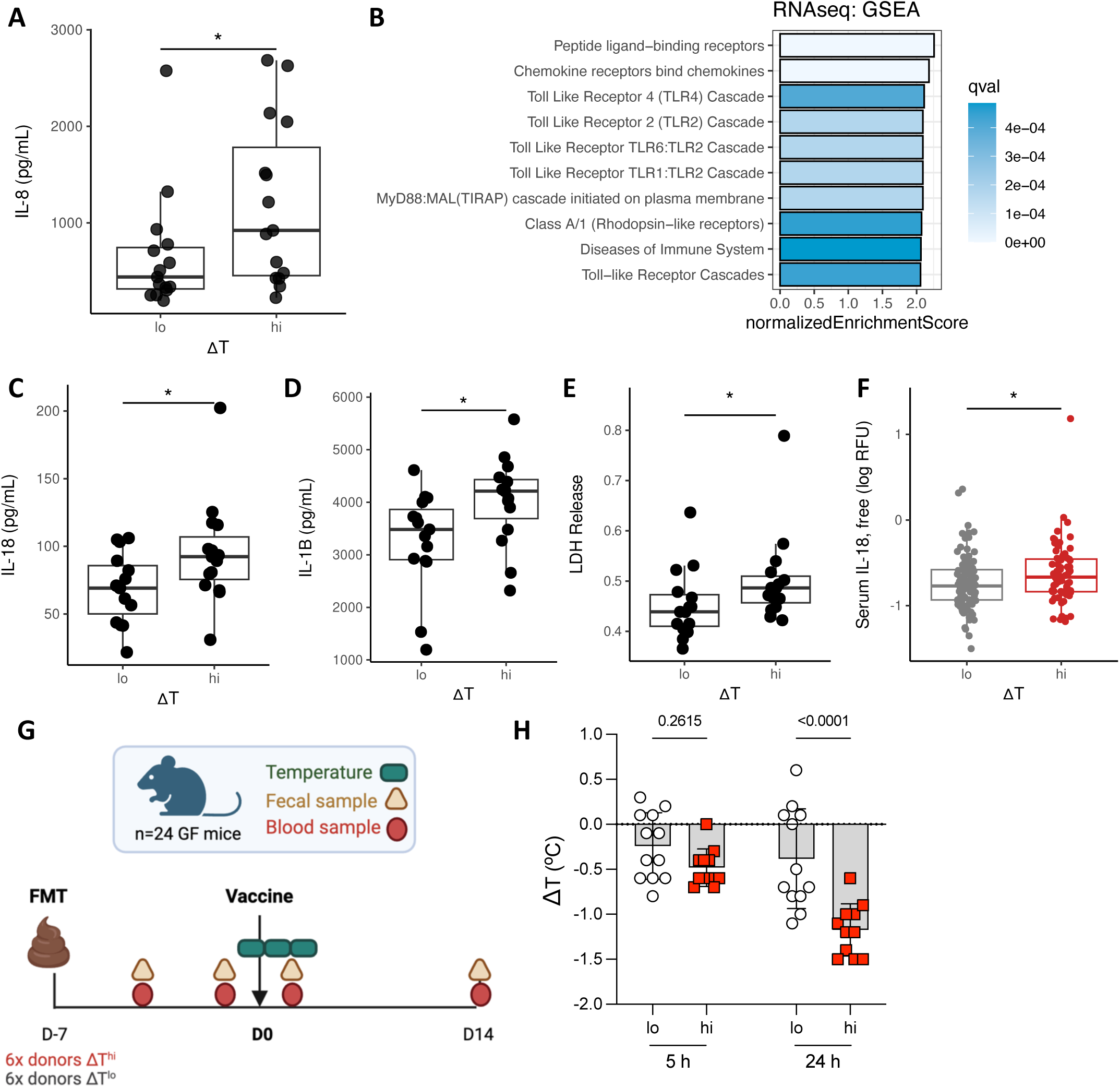
The ΔT^hi^ microbiome drives flagellin-dependent inflammation. (A) IL-8 measured in the supernatants of human colonic organoids after stimulation with fecal bacterial products derived from ΔT^lo^ and ΔT^hi^ participant samples (N=15 per group; each data point represents cells stimulated with a sample from a different person). (B) Gene-Set Enrichment Analysis (GSEA) of pathways significantly overrepresented (q<0.05) at the transcriptional level in organoids stimulated with ΔT^hi^ samples. (C-E) Human PBMCs were stimulated with fecal bacterial products derived from ΔT^lo^ and ΔT^hi^ participant samples, and in the cell supernatants (C) IL-18, (D) IL-1β and (E) cell death, via the proxy of lactate dehydrogenase (LDH) release, were measured. (F) Free IL-18 in participant serum at D-7, measured in the aptamer-based proteomics screen, and defined as the ratio of IL-18 to IL-18 binding protein. (G) Schematic for the germ-free (GF) mouse experiment. FMT, fecal microbiota transplant. (H) Change in murine body temperature (ΔT) at 5h and 24h post-vaccine. Statistical significance was determined via Wilcoxon Rank Sum test (A, C-F) and via two-way ANOVA with post-hoc Sidak’s test (Fig 5H).

In addition, PBMCs displayed significantly more cell death, IL-1β and IL-18 release, indicative of inflammasome activation, when stimulated with ΔT^hi^ compared to ΔT^lo^ fecal supernatants (Fig 5 C-E). Notably, ΔT^hi^ participants also had significantly higher levels of free IL-18 in baseline serum, indicating that basal inflammasome activity is indeed higher in these individuals (Fig 5F). Together these data suggest that products of the gut microbiome could directly drive inflammation in ΔT^hi^ individuals, likely via flagellin-mediated activation of the TLR5 and NLRC4 receptors.

Although flagellin is classically seen as a strong stimulator of innate immunity, many intestinal bacteria produce ‘silent’ flagellins with poor capacity to trigger inflammation^31^. To further clarify the role of *Waltera* flagellin in driving inflammation, we sheared flagella from pure cultures of *W. intestinalis*. Flagellin-containing bacterial supernatants activated a strong TLR5 response in TLR5 reporter cell lines (Fig S4A) and were sufficient to induce robust IL-8 responses in human organoids and IL-1β in PBMCS (Fig S4B-C). Thus, flagellins produced by *W. intestinalis* appear sufficient to explain the high inflammatory capacity of ΔT^hi^ microbiomes.

To determine if microbiome-driven inflammation would be sufficient to alter vaccine responses, we conducted a fecal microbiota transplant (FMT) experiment in mice (Fig 5G). Mice colonized with ΔT^hi^ fecal samples showed a significantly greater change in body temperature following vaccination against SARS-CoV-2 (Fig 5H) and higher levels of inflammatory serum cytokines post-vaccine (Fig S4 D-E). In summary, the fecal microbiome of ΔT^hi^ vaccine responders drives host inflammation and causes an exacerbated innate immune response to vaccination.

### Waltera growth and motility is affected by diet and controllable via a dietary additive

Diet is a powerful tool for manipulation of the microbiome. One of the microbial genes significantly upregulated with flagellins was *CsrA*, a master translational regulator that is known from bacterial pathogens to control flagellar motility in response to the nutritional environment^32^ (Fig 4A, Fig S3D). As noted previously, ΔT itself was also associated with habitual diet (Fig 1I). We therefore hypothesized that diet might drive microbiome patterns in ΔT^hi^ individuals.

To gain more detailed insight into dietary patterns, we leveraged MEDI (Metagenomic Estimation of Dietary Intake from Stool), a recently developed tool that identifies food-derived DNA present in stool^33^. MEDI confirmed differences by self-reported diet, indicating that vegetarians consumed more plants (Fig 6A), no pork, and more flax compared to omnivores (Fig S5A-B). Interestingly, several of the cytokines elevated in ΔT^hi^ participants, including pro-inflammatory IL-18, correlated negatively with the detection of plant-derived foods in stool (Fig 6B, Fig S5C), indicating a link between diet and baseline immune profiles. Nevertheless, we did not observe any direct differences in MEDI-detected foods or nutrients by ΔT category.

**Fig 6.**
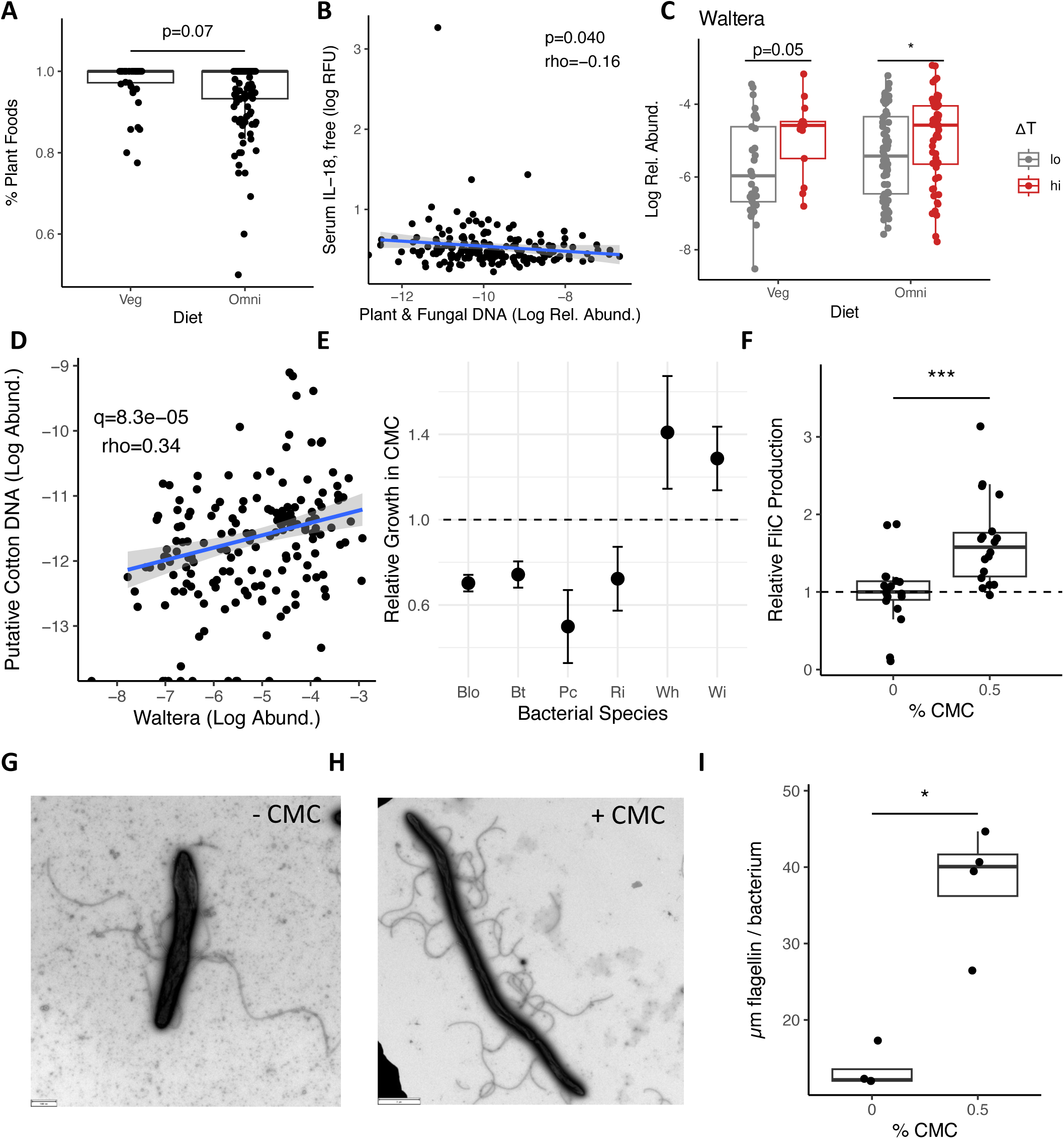
*Waltera* flagellin production is stimulated by the emulsifier carboxymethylcellulose. (A) Proportion of MEDI-detected foods which were plant-or fungi-based, across all fecal samples from each participant, by dietary category. (B) Relative abundance of all plant– and fungal-derived food DNA in fecal samples, on average across all samples per participant, compared to the detection of free IL-18 at T-7 in participant serum. (C) Log relative abundance of the genus *Waltera* by ΔT and dietary categories. Each data point represents the average abundance per person across all six samples. (D) Association between *Waltera* and MEDI-detected putative cotton DNA (assigned to *Hibiscus* genus) in stool. Average abundances per participant are plotted. (E) Relative growth of intestinal commensal bacteria at 24h in media containing 0.5% CMC compared to 0% CMC. Blo, *Blautia obeum;* Bt, *Bacteroides thetaiotamicron;* Pc, *Prevotella (Segatella) copri;* Ri, *Roseburia intestinalis;* Wh, *Waltera hominis;* Wi, *W. intestinalis.* (F) Units of FliC, detected via TLR5 reporter assay, in flagellin sheared from *W. intestinalis* cultures after growth in 0 or 0.5% CMC. (G-H) Negative stain microscopy of *W. intestinalis* grown in the absence (G, 0%) or presence (H, 0.5%) of CMC. (I) Quantification of flagellin length per cell in negative stain microscopy. Statistical significance was determined via Wilcoxon Rank Sum test (A, C, F, I) or Spearman’s correlation (B, D). In E, mean and standard error bars of three independent biological replicates are shown. In F and I, each data point represents an individual bacterial culture.

*Waltera* trended higher in omnivores compared to vegetarians (p=0.1 by mixed linear model, Fig S5D), but remained higher by ΔT even when accounting for habitual diet (Fig 6A), as did the expression of the *CsrA* gene (Fig S3D); thus, dietary category alone failed to account for microbiome differences by ΔT. Instead, *Waltera* abundance was strongly associated with the abundance of the food genus *Hibiscus* in participant stool (Fig 6D, Table S12). *Hibiscus* is prevalently detected by MEDI and associates with the consumption of certain ultraprocessed foods, particularly processed meats; it is thought to be a mis-annotation of cotton, which is a source of emulsifying agents such as carboxymethylcellulose (CMC)^33^. This led us to speculate that components of ultraprocessed foods might drive inflammatory microbiome phenotypes.

CMC has been previously shown to upregulate microbiome expression of flagellin, driving low-grade inflammation, consistent with the patterns observed in our cohort^34,35^. During *in vitro* culture, we found that CMC enhanced the growth of both *W. intestinalis* and *W. hominis*, while suppressing the growth of other common gut commensals from the *Firmicutes* and *Bacteroidetes* phyla (Fig. 6E). Moreover, growth in CMC significantly increased the production of TLR5-stimulatory flagellin by *W. intestinalis,* after normalizing for bacterial density (Fig. 6F). Individual bacterial cells produced visibly more flagella when grown in CMC, resulting in a greater total flagellar length per bacterium quantified via microscopy (Fig 6 G-I).

In summary, the dietary emulsifier CMC can effectively boost both growth and flagellin production in *W. intestinalis.* Given the impact of *Waltera* flagellins on host immunity, this may provide a tool for deliberate manipulation of the microbiome-immune axis.

## DISCUSSION AND CONCLUSION

Despite significant interest in the ability of the microbiome to modulate vaccine responsiveness, human studies to date have been largely inconclusive. Several studies have found associations between baseline immune signatures and human vaccine responses, but factors responsible for such baseline immune signatures remain undefined^36^. Here, we find that the gut microbiome stimulates vaccine reactogenicity in healthy adults via innate immune signalling. Notably, increased body temperature after vaccination associates with improved antigen-specific antibody titres, in our own study and others; however, we would have missed this microbiome signature had we focused on antibodies as the primary study outcome, in part because antibody titres were already high in this population and were driven by prior exposure. This insight may help to explain the lack of consistent microbiome-vaccine associations found in human studies to date, and indicates that the human intestinal microbiome can indeed act as a natural adjuvant for vaccination.

Importantly, we also find that vaccine reactogenicity was associated with diet: vegans and vegetarians experienced less vaccine-induced thermogenesis, and ΔT^hi^ participants had less healthy cholesterol profiles. The positive impact of plant-based diets on inflammatory and metabolic markers is well documented, and vegetarianism has also been associated with protection against symptomatic COVID-19 disease^37–40^. Our data suggests that nutrition and metabolic health might also be key factors underlying variation in vaccine side effects.

The flagellated genus *Waltera*, which was elevated in ΔT^hi^ participants, has only recently been described and isolated^41,42^; thus, much remains unknown about its basic biology and its impact on human health. However, the related genus *Acetatifactor* (from which *Waltera* was differentiated) has been associated with obesity and inflammation in murine models^43,44^. Our work indicates that *Waltera* expresses high levels of flagellin in the human gut and has the potential to be inflammatory. Moreover, we found that the dietary emulsifier CMC boosted the growth and flagellin production of *W. intestinalis in vitro*, providing a possible tool for manipulation of a microbiome-immune axis. CMC has been previously described to increase flagellin expression of the gut microbiome and drive low-level chronic inflammation in mice^34,35,45^, and large-scale studies are underway to determine its impact in humans.

Inflammation is a double-edged sword. On the one hand, inflammation promotes adaptive immune responses; encouraging microbiome flagellin expression via diet might therefore have utility for vaccination strategies in underresponsive populations. In fact, high levels of intestinal calprotectin, IL-18 and IL-8 – markers that were associated with ΔT and flagellin in our study – have been associated with better protection against COVID-19 disease itself^46^. Flagellin has already received interest for its potential as a vaccine adjuvant, and recombinant flagellin vaccines show acceptable safety profiles in humans^47–49^. Although future work is needed to test this hypothesis, native expression of flagellin by the microbiome might be particularly helpful in promoting immune responses at mucosal sites, which are typically more challenging to adjuvate^50^.

On the other hand, chronic inflammation drives pathology. Notably, multiple chronic inflammatory diseases, including inflammatory bowel disease and chronic fatigue syndrome, are associated with increases in blood antibodies against *Lachnospiraceae* flagellins^51–53^. We did not observe increases in flagellin-specific antibodies with ΔT (Fig S5 E-F), nor did the levels of intestinal inflammation measured in participants attain levels indicative of any clinical disease. Indeed, all participants were explicitly healthy young adults without co-morbidities. Nevertheless, it will be important in future work to test whether low-level, flagellin-dependent inflammation could be an early signature of disease risk. Recent epidemiological work has associated CMC consumption with increased risk of heart disease, indicating that dietary additives with microbiome effects can have long-term health consequences^54^.

In conclusion, flagellin production by the human gut microbiome drives host inflammation, promotes vaccine-induced fever, and can be manipulated via diet. This work has implications for optimizing vaccine reactogenicity and immunogenicity across diverse populations, and furthers our fundamental understanding of the role of the gut microbiome in human immunity.

## ACKNOWLEDGEMENTS

We are extremely grateful to the volunteers who chose to participate in our study during the COVID-19 pandemic, and to the members of the Clinical Trial Platform at the Institute of Tropical Medicine who supported the administration of the study, especially Anke Krauser and Eike Roggendorf. Our thanks to the Clavel lab for kindly sharing isolates of *Waltera* spp. for experimental work. We appreciated stimulating discussions on the µHEAT project with all members of the Ley lab. Funding for this work was provided by the Max Planck Society, by the Cluster of Excellence EXC 2124 Controlling Microbes to Fight Infections (CMFI), and by an EMBO Fellowship and CMFI Young Investigator Award to KH.

## Author Contributions

KEH conceived the project, performed experiments and analyses, and wrote the paper. The µHEAT Clinical Study Group implemented the human study, organized participant visits and collected samples. YHT, RK, EA, HA, DLV, SD, and AP performed experiments. NY and AT contributed to data analysis. BC, JP, PK, SK, LM, and AG provided input, resources and supervision, and edited the manuscript. ME supervised the µHEAT Clinical Study Group, contributed to study conception, and edited the manuscript. REL conceived and supervised the project and wrote and edited the paper.

## EXTENDED METHODS

### Ethics Statement

The µHEAT (<u>Micro</u>bial-<u>H</u>uman <u>E</u>cology <u>A</u>nd <u>T</u>emperature) Study was approved by the Ethics Committee of the Universitätsklinikum Tübingen under project number 398/2021BO2. All participants provided written informed consent.

Animal experiments were performed at Georgia State University according to protocol A24001.

### µHEAT Inclusion and Exclusion Criteria and Sample Size

This study targeted healthy young adults in Germany, aged 18-40, who intended to be vaccinated with a licensed vaccine against COVID-19. Detailed inclusion criteria were as follows: adults 18-40 years of age, with no underlying health conditions, belonging to the Robert Koch priority group 4 (de-prioritized for vaccination). Participants needed to want to be vaccinated against COVID-19 and be able to receive their vaccination during the study timeline. Any of the COVID-19 vaccinations approved by the Robert Koch Institute at the time of study participation (e.g., Pfizer-BioNTech, Moderna, Johnson & Johnson or other commercially available vaccines) were permitted for inclusion in the study, including booster shots. Participants needed to be able to read and understand German or English to provide informed consent for themselves. Detailed exclusion criteria were as follows: known gastrointestinal disorders (e.g., Inflammatory Bowel Disease or Celiac Disease), recent antibiotics use (any antibiotics taken in the 3 months prior to study participation), and pregnancy, as well as any health factor which placed someone at higher risk of COVID-19 disease, and therefore in a higher vaccine priority group (1-3) according to the Robert Koch Institute. Immunocompromised and immunosuppressed people were also explicitly excluded.

To estimate the ideal sample size at the outset of the study, we considered two hypotheses: first, that microbiome might influence ΔT; and second, that ΔT might affect the microbiome. For the first hypothesis, we used a power calculation based on a general linear model approach, where a continuous immune outcome (ΔT) was taken as the dependent variable and microbiota parameters (alpha diversity) were taken as the independent variable. We estimated that a biologically relevant effect size (f2) would be 5% of the variation in immune responsiveness explained by baseline microbiota diversity. We used a significance level (α) of 0.05 and a power of 0.90 and implemented the calculation in R using the pwr.f2.test function. This yielded a sample size of 256 participants. To further confirm that this sample size was sufficient for the second hypothesis, we estimated that at least 20% of participants (18-40 years old) in our study would experience ΔT, or 51 out of

256 participants. We used a power calculation based on a two-sample t-test to compare microbiome parameters of participants with and without ΔT. Based on data previously collected by our group on the microbiomes of healthy adults in Tübingen, we estimated an effect size of 4.6 units Shannon Diversity and a standard deviation of 0.68. Taking uneven sample sizes of n1=51 (with ΔT) and n2=205 (without ΔT), and α=0.05, a power calculation with pwr.t2n.test suggested we would be highly powered (99%) to also detect differences in the microbiome induced by categorical ΔT in this study design.

A total of 182 people completed participation between November 2021 and May 2022, at which point the majority of people in Tübingen who wished to be were already fully vaccinated and therefore recruitment was ended.

We further excluded participants from downstream analysis who (a) experienced a COVID-19 or other infection during the study; (b) provided unreliable thermometer data, such that estimating ΔT was not possible; or (c) received a non-mRNA-based vaccine (n=1 who received Novavax, an adjuvated vaccine). This yielded a final sample size for analysis of 171 people. Final metadata for participants can be found in Table S1.

Within this population, n=2 (1.7%) received dose 1, n=20 (11.7%) received dose 2, n=143 (83.6%) received dose 3 and n=6 (3.5%) received dose 4 during the study. N=157 (91.8%) of doses were BioNTech and the remaining N=14 (8.9%) were Moderna.

### µHEAT Study Design

Study participation for each subject began 1 week (6-9 days) prior to their vaccine (Fig 1A). At an enrollment visit, participants provided a blood sample and filled out a demographic questionnaire which included information on prior vaccination, prior COVID-19 infection, as well as lifestyle factors such as diet, smoking and alcohol consumption. Height and weight were measured and oral temperature was taken under the supervision of a study physician. Participants were provided with a study thermometer to take home and instructed by a study physician how to properly take oral body temperature. Participants were also provided with a commercial stool collection kit (Zymo DNA/RNA Shield) to use for at-home collection, as well as an extra empty 10 mL tube (no stabilization solution) to be filled in the 24h prior to their last clinic visit days for immediate freezing at the clinic.

Participants took their oral body temperature 3 times per day and recorded it in their study diary, along with the time of measurement. They were instructed to take their temperature at approximately the same times every day: once each morning, afternoon and evening. The study diary also provided spaces to record other possible systemic symptoms of vaccination every day (before and after the vaccine), namely headache, fatigue, chills, loss of appetite, diarrhea, and fever (subjective / self-reported). Participants were also asked to record any instance of pain-or fever-relieving medication they took throughout the study.

For each stool sample collected at home, participants also recorded in their diary the date and time, and estimated the Bristol stool score according to a provided Bristol stool score chart. They were instructed to provide 3 samples in the week before vaccination and 3 samples in the week after vaccination.

At a final study visit two weeks after vaccination, participants returned their study diaries and fecal kits and provided a final blood sample for antibody titres. Freshly provided fecal samples (sample #7, no buffer, collected within prior 24h) and serum were immediately frozen at –80°C. Buffered fecal samples (samples #1-6) were transferred to the laboratory at MPI Tübingen where they were aliquoted for DNA and RNA extraction prior to freezing at –70°C.

### Temperature

To define ΔT, we first defined a personal baseline temperature per person as the median body temperature for a given time of day (morning, afternoon, or evening recording) for that person, in the week prior to vaccination. ΔT was then considered as the maximum observed increase in body temperature relative to baseline for the same time of day, in the week following vaccination. This variable ranges between 0.2°C and 2.4°C with a median of 0.7°C and a mean of 0.8°C. We categorized people as ΔT^hi^ or ^lo^ based on the group mean. In practise, participants were therefore considered as ‘ΔT^hi^’ if they had an increase in body temperature following vaccination of 0.8°C or more above their own personal baseline. For comparison, the average standard deviation of personal body temperature at baseline was 0.3°C.

For self-reported prior fevers, participants were asked if they had experienced a fever (body temperature ≥38.0°C and/or fever symptoms such as aches and chills) to prior doses of the SARS-CoV-2 vaccine or at any point during the prior two years.

### Diet

Self-reported vegans and vegetarians were designated as following a plant-based diet. All other self-reported dietary categories, including flexitarians, pescatarians, ketogenic and fish-free diets, were considered omnivores unless otherwise specified.

### Serum antibodies

Anti-SARS-CoV-2 IgG was quantified in serum via ELISA using the Invitrogen Human SARS-CoV-2 Spike (trimer) IgG ELISA kit, according to the manufacturer’s instructions. Serum samples were diluted 1:25 000 in kit assay buffer and were run in a randomized order.

### Serum metabolomics

For metabolomics, 45 µL of serum was extracted using 244 µL of a butanol-methanol solvent (1:1 butanol:methanol containing 1.13 µg/mL EquiSPLASH and 0.32 µg/mL Trp_13_C_11_). Samples with solvent were vortexed for 15 minutes at room temperature using a multi-sample vortex, centrifuged 13000g for 10 min, and the supernatants were transferred to clean Eppendorf tubes, aliquoted and stored at –20°C until mass spectrometry analysis. Serum was processed and run in a randomized order. A QC pool was created by mixing 5 µL of each sample; an aliquot of this QC pool was then extracted every 10 samples to account for temporal variation in mass spec data.

Samples were analyzed in a Dionex UltiMate 3000 HPLC (Thermo Scientific, USA) coupled with a high-resolution mass spectrometer (Impact II mass spectrometer, Bruker, Germany). HPLC separation and mass spectrometric conditions were as reported previously^55^. Acquired data were then converted to mzML format using MSConvert tool from ProteoWizard^56^ prior to data processing by Mzmine3^57^. Features were annotated using MoNA database and/or manually checked.

Preprocessed metabolomics data was further blank-filtered at a cutoff of 0.3, such that metabolites more than one-third as abundant in blank samples compared to real samples were excluded. Analysis was then performed separately on negative– and positive-mode data using MaAsLin2 linear models. Data was normalized via total sum scaling and log transformation and a prevalence cutoff of 0.25 was applied. Age, sex, and habitual diet were taken as covariates and p-values were corrected via FDR.

### Cholesterol assay

Serum cholesterol was quantified using the HDL and LDL/VLDL Cholesterol Assay Kit (Antibodies Online ABIN2345055) according to the manufacturer’s protocol. Serum samples were run at a 1:4 total dilution in a randomized order.

### Serum proteomics

Serum samples were profiled by Somalogic aptamer technology for a semi-targeted panel of 1500 immune and metabolic proteins. The custom panel can be found in Table S1. Serum order was randomized for profiling and per-plate control normalization steps were performed by Somalogic. Analysis was then performed using either Wilcoxon Rank-sum tests or MaAsLin2 linear models correcting for age, sex, habitual diet and BMI. All p-values were corrected via FDR.

### Fecal calprotectin

Fecal calprotectin was measured in fresh-frozen fecal samples via the human Calprotectin L1/S100-A8/A9 Complex ELISA Kit (Invitrogen EH62RB) according to manufacturer’s instructions. Fecal supernatants were first generated for all samples according to the Methods section ‘generation of fecal supernatants’. Samples were run at 1:20 dilution in a randomized order using an overnight sample incubation step. For detection of calprotectin via RT-qPCR in ZYMO Shield-buffered samples, RNA was extracted using the ZymoBIOMICS RNA Miniprep Kit, including the DNAse step and eluting in 85 µL. cDNA was generated using RevertAid First Strand cDNA Synthesis Kit (ThermoFisher), with oligo dT primers and an incubation of 60 min at 42°C plus 5 min at 70° in a thermocycler, from RNA normalized to 20 ng/µL. qPCR was then performed using Platinum SYBR Green Master Mix for a total reaction volume of 10 µL with 2 µL of cDNA and 0.4 µL of primers. Primers were as follows: S100A9_F, GCACCCAGACACCCTGAACCA; S100A9_R, TGTGTCCAGGTCCTCCATGATG; GAPDH_F, GGAGCGAGATCCCTCCAAAAT; GAPDH_R, GGCTGTTGTCATACTTCTCATGG (Harvard Primer Bank). Cycling conditions were: 50°C for 2 min, followed by 40 cycles of 95°C for 15 sec, 55°C for 1 min. Undetectable CT values (no amplification) were imputed as 40 cycles. Fold changes were calculated using the formula 2^-ΔΔCT^, in which ΔCT compared S100A9 to GAPDH per sample and ΔΔCT was relative to the average ΔCT for all samples in that cDNA batch (plate). Samples in which GAPDH was completely undetectable (both technical replicates CT value ≥40) or in which the calculated fold change exceeded the higher and lower quartiles of the total fold-change distribution by more than 10x (i.e., fold changes >100 or <0.000001, which were suspected to reflect technical errors) were excluded from further analysis.

### Metagenomic sequencing

A total of 1145 longitudinal samples were processed for DNA extraction and metagenomics sequencing. DNA was extracted in a randomized order using ZymoBIOMICS 96 MagBead DNA kit. Samples were first bead-beat using Zymo “bashing” bead tubes with 8 cycles of 40 seconds, 6 m/s and 5 minutes rest between cycles on a BioSpec Bead Beater. Samples were then transferred to plate format and processed on a Hamilton robot. Each plate contained at least one extraction blank and one mock microbial community (Zymo D6300) for quality control.

Library preparation on the extracted DNA was performed as previously described^58^, using a Nextera (Illumina, San Diego, USA) Tn5 tagmentation reaction to fragment and ligate adaptors in a single reaction. Following a cleanup step, DNA was amplified via Q5 polymerase in a total volume of 25 µL over 8 PCR cycles, using multiplexed barcoded primers. Consistent amplification was confirmed via qPCR. Samples were then pooled at equal volumes. A final PCR cleanup followed by size selection (300-800 bp) with BluePippin (Sage Science, Beverly, USA) on a 1.5% gel was performed, and final library size and distribution was confirmed on a Bioanalyzer using a High-Sensitivity DNA Chip. Sequencing was performed by Novogene on a NovaSeq6000 using a P4/300 cycle kit in three randomized batches.

Sequencing data was processed using in-house standardized Snakemake-based pipelines. A quality-control pipeline used bbduk to remove Illumina adapters and bbmap to filter out reads mapping to the human genome (https://jgi.doe.gov/data-and-tools/bbtools/). Seqkit^59^ and fastqc (https://github.com/s-andrews/FastQC) were applied before and after QC for inspection read depth and quality. Post-QC samples had an average read depth of 12.7 M. Four samples with a depth of less than 0.5 M, comparable to sequenced blank controls, were excluded from further analysis. In total, after technical and biological exclusions, 1022 samples with sequencing information from 171 participants were retained for analysis.

Diversity and abundance profiling was performed using kraken2^60^ and bracken^61^ for abundance estimates and for alpha and beta diversity, leveraging a custom database built via Struo2^62^ on GTDB release 207^63^. Diversity was performed on data rarefied to 5 M reads, which retained 942 samples. Humann3^64^ was used for gene-level abundance estimates.

CoPTR (https://coptr.readthedocs.io/en/latest/index.html) was used for estimation of bacterial growth rates. inStrain^65^ was used to detect strain-level persistence and evolution. MEDI^33^ was applied for classification of food-derived DNA.

Abundance differences were tested using MaAsLin2^30^ linear models on log-scaled longitudinal relative abundance data, with an abundance cut-off of 0.0001 and a prevalence cut-off of 0.25. Sequencing depth, Bristol Stool Score and the amount of time the sample spent at room temperature before freezing were taken as technical covariates and participant ID was treated as a random effect. P-values were corrected via FDR.

### Transcriptomic sequencing

Samples were manually selected for RNAseq based on the presence of complete and clear ΔT profiles, attempting to maximize ΔT differences by category (i.e. highest and lowest ΔTs in the study population) while matching as closely as possible between high– and low-ΔT samples for age, sex and vaccine history. Samples used for RNA-seq are flagged in Table S1. This subpopulation was overall similar to the total population, with basic demographics as follows: age 24.5 yrs, 60.5% female, 84.2% dose 3, and 89.5% BioNTech.

Similar to metagenomic library preparation (above), samples were first bead-beat with Zymo bead bashing tubes as described, but with reduced program of 4 bead-beating cycles, 40 sec, 6 m/sec with 1 min cooling on ice between samples. RNA extraction then proceeded with either the ZYMO RNA miniprep kit (pool 1) or with the NEB Monarch tRNA Miniprep Kit (pools 2-3), which have similar extraction chemistries. In both cases, an optional reduced ethanol volume was used at the initial ethanol step to exclude small RNAs ≤200 bp. RNA quality was spot-check via Bioanalyzer and quantified via Qubit. Library preparation was then performed using an Illumina Stranded Total RNA Prep, Ligation with Ribo-Zero Plus Microbiome. Library quality was checked on the Bioanalyzer and quantified via Qubit, normalized and pooled equimolar. A final 0.9x bead clean was performed on the final pool prior to sequencing.

Sequencing was performed by Novogene on a NovaSeq6000 with a P4/300 cycle kit in three randomized batches.

Microbial transcriptomic sequencing data was preprocessed and profiled using the same pipelines as for metagenomic sequencing, above. Post-QC samples had an average read depth of 49.2 M. Diversity analyses were performed on data rarefied to 5 M reads, which retained 226 samples.

Gene-level abundance differences in transcriptomic data were tested using MaAsLin2 linear models on log-scaled longitudinal relative abundance tables, with an abundance cut-off of 0.00001 and a prevalence cut-off of 0.25. Sequencing depth, Bristol Stool Score and the amount of time the sample spent at room temperature before freezing were taken as covariates and participant ID was treated as a random effect. P-values were corrected via FDR.

### *In vitro* bacterial temperature phenotypes

High-throughput bacterial growth curves were performed as previously described^58,66^. Briefly, human gut commensals (Table S2) were grown anaerobically in mGAM media. Strains were inoculated from glycerol at 37°C, subcultured overnight 1:100 at 37°C, and finally inoculated into 100 µL cultures in 96 well plates at a normalized O.D. of 0.02 under the desired experimental temperature. Due to the necessity of changing the incubator temperature, each temperature condition was run separately. Half of a plate (48 technical replicates) per strain per condition were used, using an automated stacker and plate reader to read O.D. over a 24 hour period, and each strain was run over 2-3 biological replicates at that temperature. The growth rate of each species was then calculated in R by manually separating the linear part of each growth curve (exponential phase) and then obtaining the slope via linear fit.

For RNA-seq of *Prevotella copri,* the bacterium was cultured and subcultured as described above in 10 mL volumes. Temperature shock was then performed for 1 hour via water baths at 32°C and 42°C, versus a control condition maintained at 37°C. RNA from each full culture was extracted in QIAZOL using the miRNEAsy kit (Qiagen) and quantified via Qubit. RNAseq library preparation was performed using the Illumina Stranded Total RNA Prep, Ligation with Ribo-Zero-Plus. Sequencing was performed in-house on a NextSeq2000. Analysis of the RNA-seq data followed an in-house custom pipeline which used BBDuk to trim adaptors (https://jgi.doe.gov/data-and-tools/bbtools/), FastQC to assess read quality (https://github.com/s-andrews/FastQC), and Bowtie2 to align reads to the *P. copri* reference genome^67^.

### Generation of fecal supernatants

Freshly frozen (i.e. unbuffered) fecal samples collected within 24 hours of the last study visit were used for protein-based assays and experimental work. Fecal supernatants were prepared by resuspending stool in PBS containing an EDTA-free protease inhibitor at a concentration of 200 mg/mL. Approximately 150 mg (+/-10%) was aliquoted from frozen stool using a biopsy punch and weighed into an empty 2 mL FastPrep tube; the exact weight was recorded, and the PBS volume was adjusted to achieve a final concentration of 200 mg/mL (approximately 750 µL). Samples were then homogenized using a 2×2 min FastPrep cycles (without the addition of beads) at 6 m/s, which should also achieve shearing of bacterial flagellins. Bacteria and debris were pelleted via two spin cycles at 13700g 10 min 4°C. For cell culture experiments, the supernatant was further sterilized and flagella were depolymerized via a 15 min boil at 98°C in a heating block. For other ELISA assays unboiled supernatant was used. Supernatants were stored at –70°C until used.

### Organoid experiments and RNAseq

Human colonic enteroids line xcol4 were thawed, passaged and seeded according to the protocol “Human Enteroid Protocol Handbook” by Julia Y. Co, Stanford University School of Medicine, as previously described^31^. Briefly, organoids were grown in 24-well-plates with 400 µL / well of IntestiCult Human Basal Medium (StemCell Technologies) supplemented with Organoid Supplement (StemCell) (“Growth Media”). For passaging, 1 mL of TrypLE Express / well was used to digest organoids, which were then aspirated and pooled into a falcon tube and incubated for several minutes at 37°C, before inactivating with FBS and spinning at 500g for 3 min to pellet cells. Cells were then resuspended in cold Matrigel on ice and seeded into a fresh 24 well plate at 40 µL / well. After passaging, media contained additional inhibitors (10 µM of Y-27632 and 250 nM of CHIR99021, BioTechne) (“Passage Media”) for 2-3 days until replaced with fresh growth medium. Cells were seeded approximately one week before the experiment until mature organoids were observed.

Fecal donors for cell culture experiments are flagged in Table S1 and were chosen by matching as evenly as possible according to age, sex and vaccination history, while maximizing differences in ΔT responsiveness. Frozen fecal supernatants were thawed on ice and then boiled for 15 minutes to sterilize and to depolymerize any flagellins. Any remaining debris was settled with a short spin. In addition, flagellin and empty vector controls (extracted as described above) were thawed on ice and diluted in PBS to a working stock concentration of 100 nM. Purified LPS controls were used at 1 µg/mL and purified calprotectin at 10 ng/mL working concentration. The organoid media was exchanged for 360 µL of fresh Growth Media. 40 µL of each fecal supernatant or control was added per well, using two wells (technical replicates) per sample. Moreover, two replicates or batches of organoid cells were used, with different participant fecal samples but identical flagellin controls per batch. From each experiment, cell supernatants were collected at 4h (with replacement of 200 µL fresh media) and at the endpoint of 18h. At 18h, RNA was also saved for extraction according to the ReliaPrep RNA Cell Miniprep System with modifications for organoid cells (Promega protocol Application Note #AN296). Briefly, after removal of media, 200 µL of cold LBA buffer containing TG was added per well and cells were resuspended via pipetting. This suspension was then frozen at –70°C until extraction.

IL-8 was measured in undiluted or 1:1.3 diluted cell supernatants using the R&D DuoSet IL-8 ELISA according to kit instructions.

RNA extraction was continued in a randomized order across both experimental replicates, according to kit instructions. RNA libraries were generated using the NEBNext Ultra II Directional RNA Library Prep Kit for Illumina in randomized batches of 24 samples. Sequencing was performed in-house on a NovaSeq2000.

### PBMC experiments

Human PBMCs were isolated from a healthy adult volunteer. Briefly, whole human blood was diluted 1:1 with Hank’s Balanced Salt Solution and separated via Ficoll-Hypaque gradient and centrifugation (25 min, 810g, no brake). Isolated PBMCs were washed 3x with HBSS (10 min, 200g), counted, and resuspended in media (RPMI, 10% FBS) at 0.5 x 10^6^ cells / mL. Cells were then seeded into a 96-well-plate at 180 µL / well, and stimulated with 20 µL of fecal supernatant for 20-24h. Cell supernatants were used to assess cell death using an LDH Release Assay (ab65393, AbCam), and IL-1β and IL-18 were measured via ELISA (R&D DuoSet ELISAs, DY201 and DY318).

### Waltera flagellin

*Waltera intestinalis* and *W. hominis* were a generous gift from the Clavel lab. *Waltera* was routinely grown in anaerobic YCFA broth without the addition of vitamins, using stoppered Balch tubes with an N_2_/CO_2_ atmosphere. Overnight cultures were inoculated using 100 µL glycerol stock aliquots into 10 mL of YCFA for 24 hours at 37°C without shaking. Subcultures were then prepared using 100 µL of 24h culture into 10 mL of fresh YCFA overnight at 37°C without shaking.

To shear flagellins from *Waltera* cultures, bacterial cells were pelleted and resuspended in PBS containing protease inhibitor, and normalized (concentrated) to an O.D. of 0.5. Cells were then Fastprepped, spun and boiled as described previously for the fecal supernatant preparation. The end product was a sterile supernatant preparation, containing neither bacterial cells nor media nor secreted metabolites, only heat-resistant bacterial products including flagellin. *Waltera* sheared flagellins were then added to organoids and PBMCs at 20 µL / well as described above.

### TLR5 reporter assay

The HEK TLR5 reporter cell line (Invivogen) was grown routinely in DMEM containing 10% FBS. For reporter assays, a single confluent T75 was used to seed a 96 well plate, using 180 µL of resuspended cells per well and directly adding 20 µL of stimulus. FliC (expressed and purified in-house as previously described, according to the genetic sequence for *Salmonella typhimurium*^31^) was used as a standard curve for TLR5 at a top concentration of 10 µg/mL. Plates were incubated at 37°C 5% CO2 for 24h. Per well, 180 µL of QuantiDetect dye was then mixed with 20 µL of cell supernatants and O.D. 635nm was read at 10 minutes and at 30 minutes.

### Flagellin serum ELISAs

Flagellin ELISAs were performed as previously described^34^. Plates were coated overnight at 4°C with 100 ng per 100 µL well of stimulatory FliC, or with an equimolar equivalent of the silent flagellin FliC-1 (UniRef50 ID R6ZZ21), or with 100 µL of a purified empty-vector control (no flagellin), all expressed and purified in-house as previously described^31^. Plates were blocked with 200 µL of PBS 1% BSA for 1h. Serum was used at 1:200 dilution in a randomized order for 2 hours. An anti-hIgG HRP (Thermo, Cat # 62-8420) was used at 1:4000 for 1 hour. ELISAs were detected, stopped and read according to standard protocol^34^.

### Waltera growth curves

Growth curves from subcultures were inoculated in an anaerobic chamber to a normalized starting O.D.600 of 0.02 in YCFA with or without 0.5% CMC, in a 96 well plate sealed with a BreatheEasy permeable membrane. O.D. was measured over a 24-48h period with double orbital shaking at 37°C. Comparison species *Bacteroides thetaiotamicron, Prevotella copri, Roseburia intestinalis* and *Blautia obeum* (Table S13) were also grown in YCFA following the same protocol as for *Waltera*.

### Waltera microscopy

Fresh subcultures of *W. intestinalis* in YCFA with or without 0.5% CMC were prepared for microscopy. For negative stain, glow-discharged formvar– and carbon-coated copper grids were incubated with concentrated cells, and stained with 1% uranyl acetate. Images were acquired with a CMOS camera (TemCam-F416, TVIPS, Gilching, Germany) mounted on a Tecnai Spirit transmission electron microscope (Thermo Fisher Scientific, Eindhoven, The Netherlands) operated at 120 kV. Flagellin length from negative stain images were quantified using ImageJ’s line function by tracing each flagellin present in the image and calculating the resultant line length in µm. Flagellin length was then normalized to the number of bacterial cells visible in the image to obtain µm flagellin / cell. Partially visible bacteria in the image were counted as 0.5 of a bacterium. Flagellin measurements were performed blinded to the experimental condition (YCFA vs CMC).

### Animal experiments

6-week-old germ-free C57BL/6N mice (Taconic) were inoculated by oral gavage with fecal slurries prepared from ΔT^hi^ and ΔT^lo^ donors. Six ΔT^hi^ and six ΔT^lo^ donors were selected, and two cohoused mice were used per donor (total N=24). Donors were split evenly by sex (3 female and 3 male per ΔT category), and sex-matched to recipient mice to ensure balanced groups. Fecal donors are flagged in Table S1 and were chosen by matching as evenly as possible according to age, sex and vaccination history, while maximizing differences in ΔT responsiveness. Mice were colonized for 1 week prior to vaccination with 5 µg of Delta/BA.5 Bivalent vaccine (mRNA SARS-CoV-2 vaccine). Body temperature at 0h, 5h and 24h post-vaccine were tracked via rectal thermometer. Blood was collected intraorbitally pre– and post-vaccine to track serum cytokine responses using ELISA kits (IL-6; DY406, Cxcl1; DY453, R&D).

### Statistical analysis

Statistical analysis was performed in R 4.1.2 and in GraphPad Prism 10. The details of each statistical test are reported in the figure legends or text and in Tables S3 and S5-12.

### Data and Code Availability

Full markdowns for all R code used in the paper are available on Github (https://github.com/leylabmpi/uHEAT). Sequencing data is uploaded to the SRA under accession numbers PRJEB85985 (human fecal metagenomes and transcriptomes) and PRJEB85993 (in vitro RNA-seq data) and will be made public upon acceptance of the paper.

## SUPPLEMENTAL FIGURES

**Fig S1.**
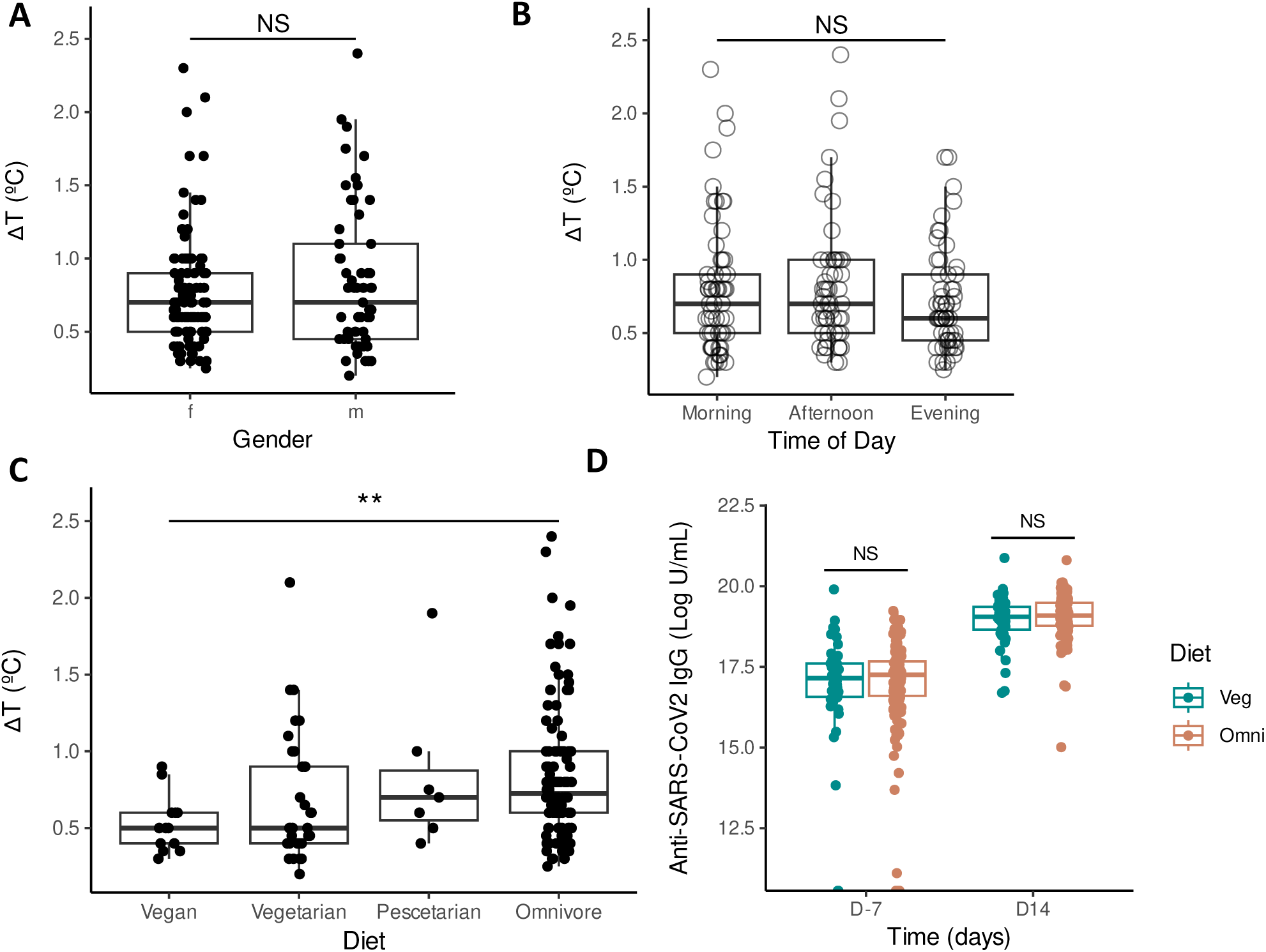
Degree of post-vaccine temperature increase (ΔT) according to (A) gender, (B) time of day at which the maximum temperature was recorded, and (C) dietary category. (D) Serum anti-SARS-CoV-2 IgG pre– and post-vaccine, according to dietary category. Statistical significance was determined via Wilcoxon Rank-Sum test (A, B, D) or Kruskal-Wallis test (C).

**Fig S2.**
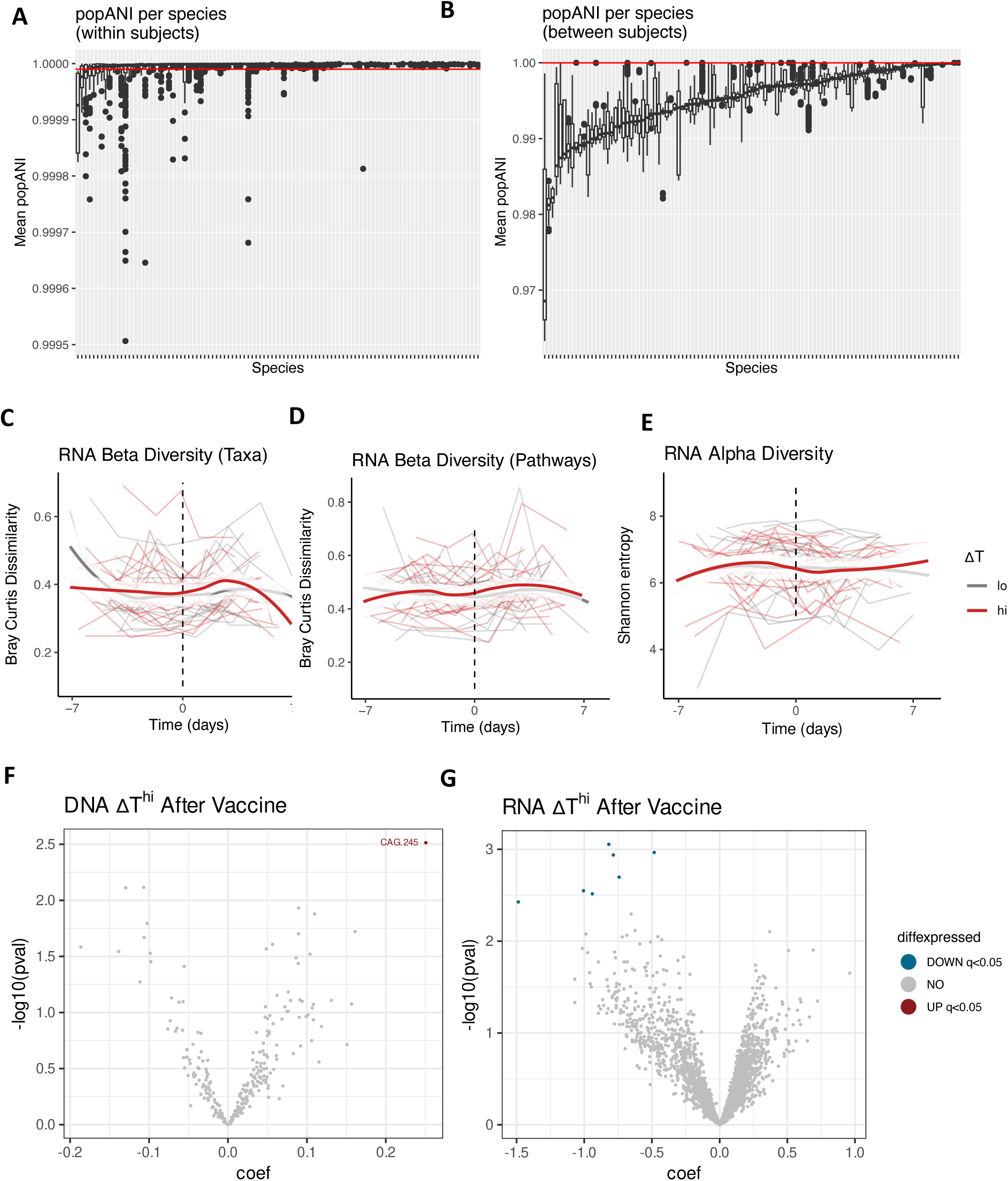
(A-B) Mean popANI per species within samples from (A) the same participant and (B) different participants. The red line shows a mean popANI of 0.99999, considered to be the cutoff for a unique strain. Species names are omitted for legibility. (C-D) Bray-Curtis dissimilarity of each sample compared to every other sample from that individual, over time, based on transcriptomic data at (C) the taxonomic level and (D) the level of gene pathways. (E) Transcriptomic alpha diversity over time. (F-G) Volcano plots of taxa significantly different in ΔT^hi^ participants after vaccination compared to other samples, based on (F) metagenomic and (G) transcriptomic data. Statistical significance was determined via MaAsLin2 linear models taking sequencing depth, Bristol Stool Score, and time at room temperature as technical covariates, treating participant ID as a random effect, and adjusting p-values via FDR (F-G).

**Fig S3.**
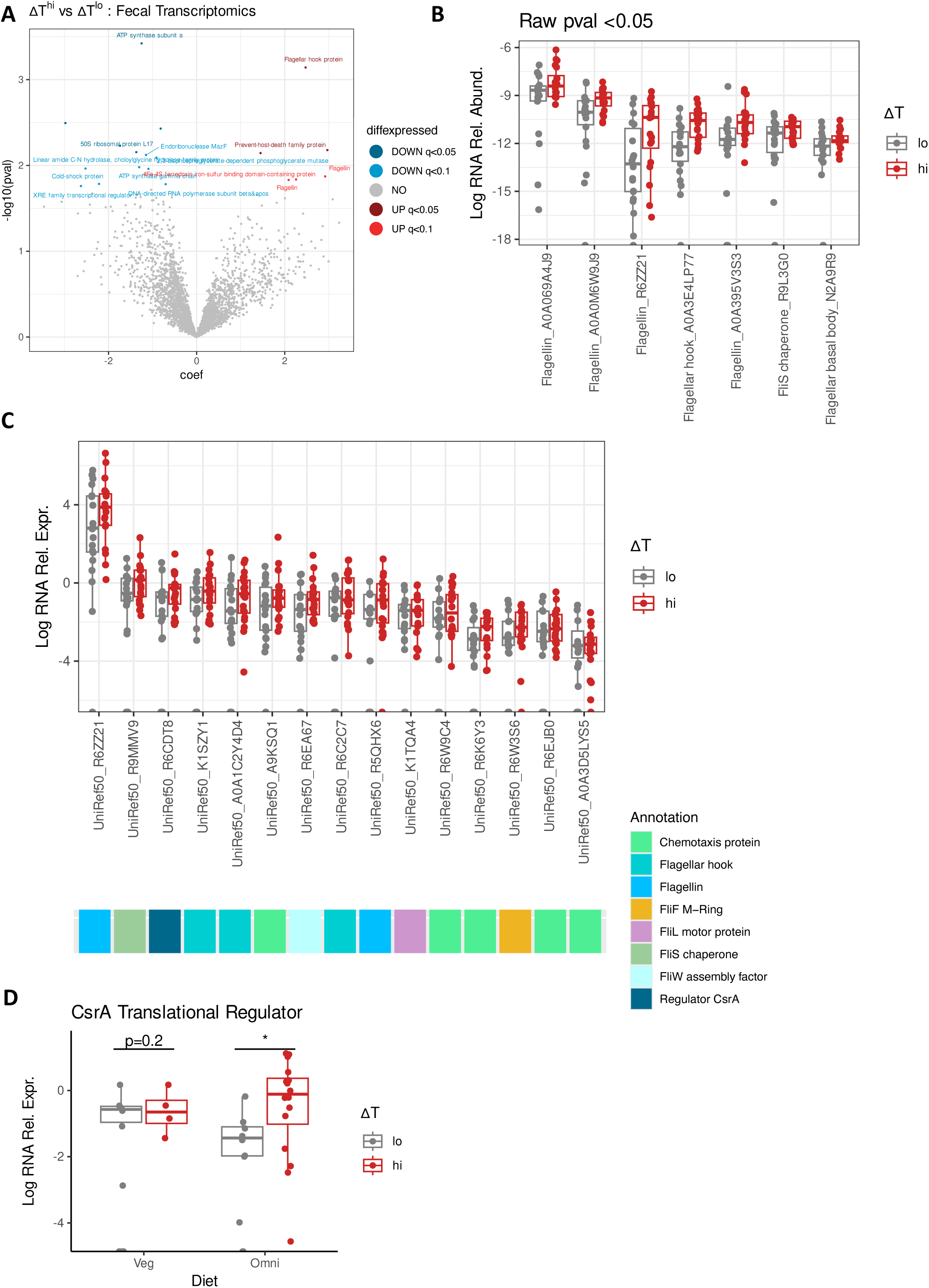
(A) Volcano plot of genes (UniRef50) significantly enriched and depleted at the transcriptional level, considering only RNA relative abundance, in ΔT^hi^ compared to ΔT^lo^ individuals. All significant genes are labelled. (B) The log-normalized transcriptional relative abundance of all genes that are significantly different by ΔT (unadjusted p<0.05) and have an annotated role in motility. (C) The log-normalized transcriptional relative expression, normalized to DNA abundance of that same gene, of all genes that are significantly different by ΔT (adjusted q<0.05) and have an annotated role in motility. (D) The log-normalized transcriptional relative abundance of CsrA by dietary and ΔT categories. Each data point represents the average abundance per person across all six samples (B-D). Statistical significance was determined via MaAsLin2 linear models taking sequencing depth, Bristol Stool Score, and time at room temperature as technical covariates and treating participant ID as a random effect (A-C), and by Wilcoxon Rank Sum test (D).

**Fig S4.**
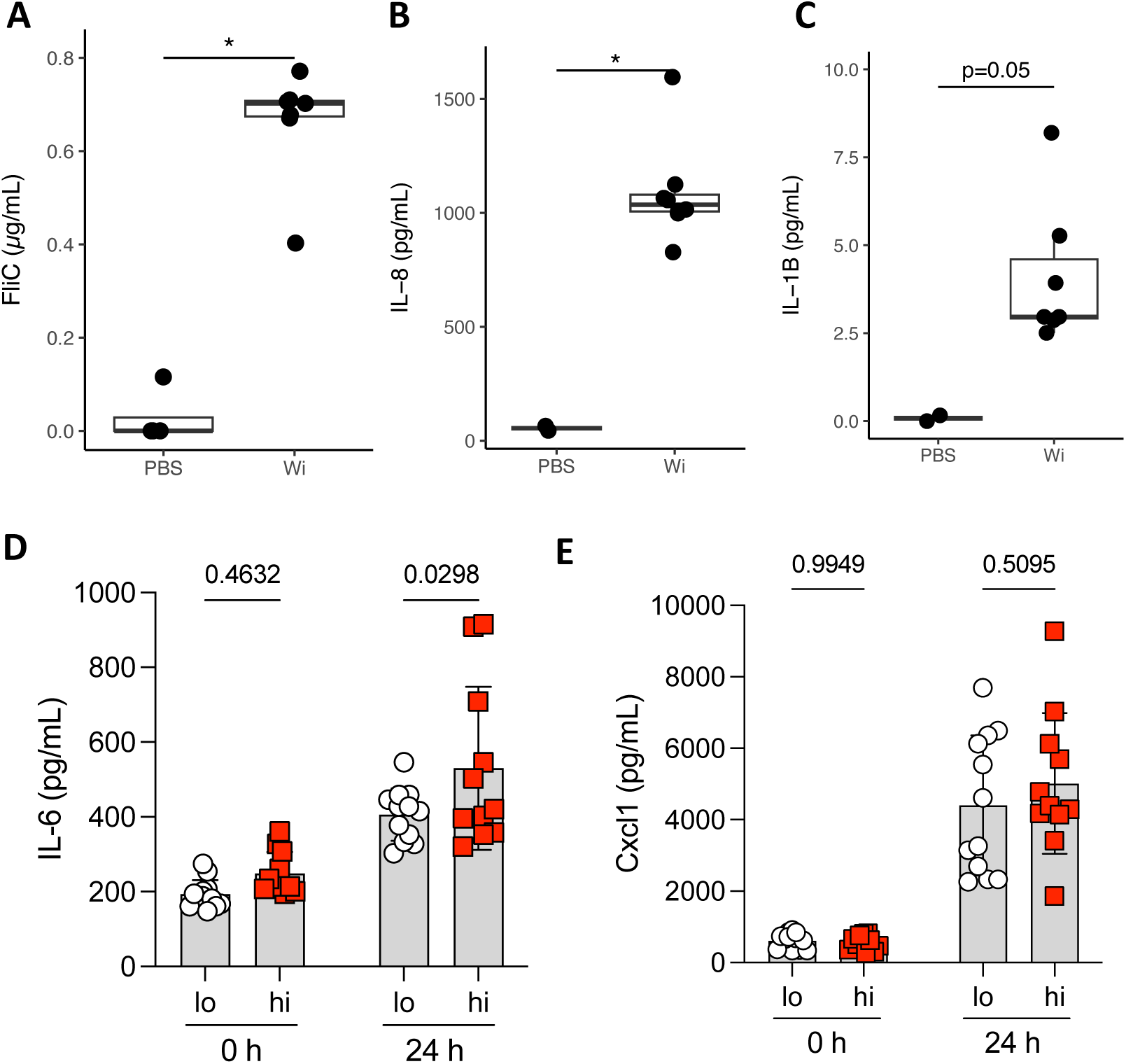
(A) Measured units of FliC equivalents in sheared flagellins from *W. intestinalis*, according to a TLR5 cell reporter assay. (B) IL-8 induced in human colonic organoids by stimulation with sheared *W. intestinalis* flagellins. (C) IL-1β induced in human PBMCs by sheared *W. intestinalis* flagellins. (D-E) Serum (D) IL-6 and (E) CXCL1 detected in gnotobiotic mice post-vaccination. Statistical significance was determined via Wilcoxon Rank Sum test (A-C) and via two-way ANOVA with post-hoc Sidak’s test (Fig S4 D-E).

**Fig S5.**
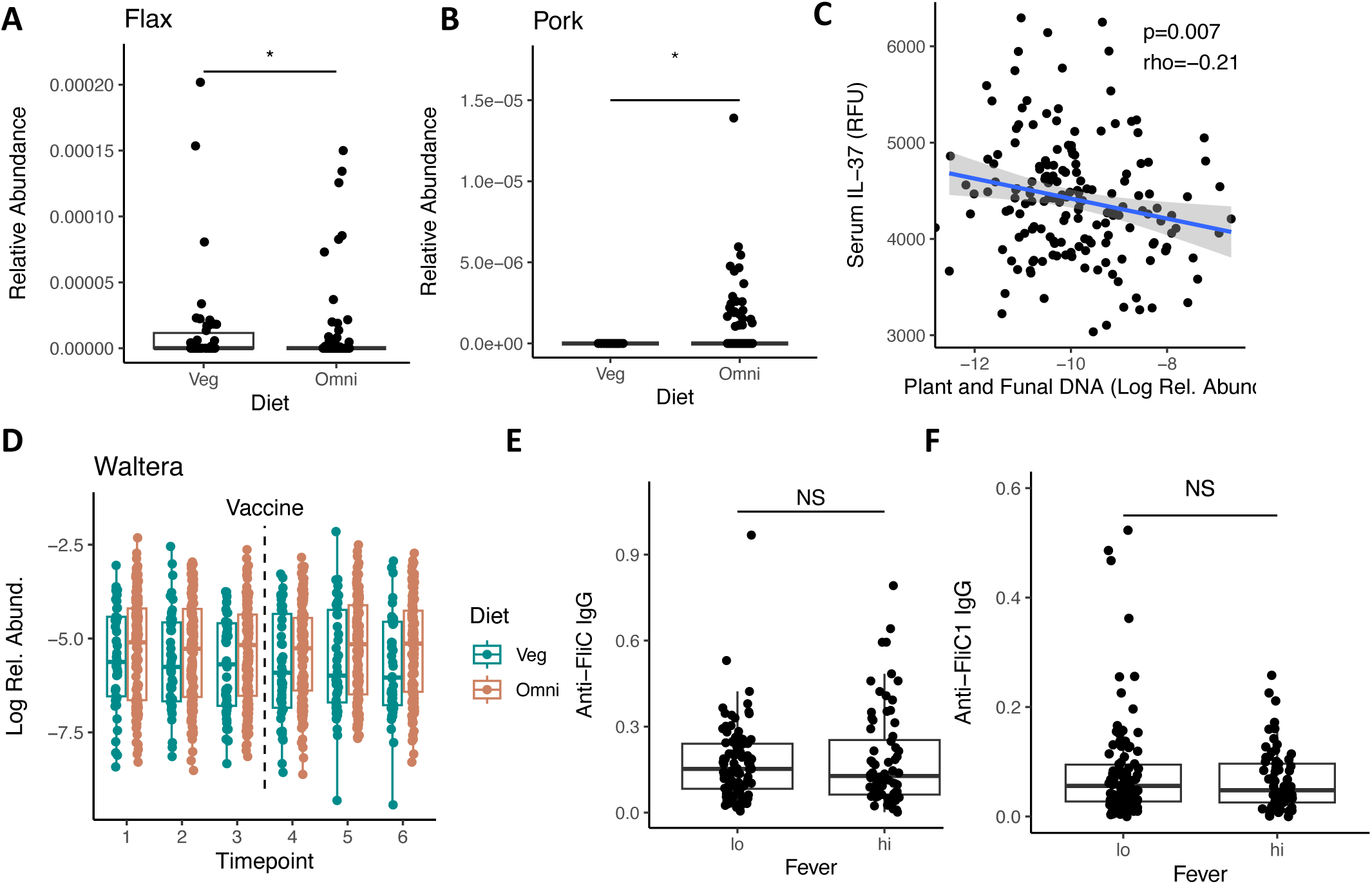
(A-B) Relative abundance of flax (A) and pork (B) detected by MEDI on average per person. (C) Relative abundance of all plant– and fungal-derived food DNA in fecal samples, on average across all samples per participant, compared to the detection of IL-37 at D-7 in participant serum. (D) Log relative abundance of the genus *Waltera* by dietary category over time. (E) Serum antibodies against purified FliC at D-7. (F) Serum antibodies against purified FliC-1, a *Lachnospiraceae* silent flagellin corresponding to UniRef50 ID R6ZZ21, at D-7. Statistical significance was determined via Wilcoxon Rank Sum test (A-B, E-F) or Spearman’s correlation (C).

## SUPPLEMENTAL TABLE LEGENDS

**Table S1.** Metadata of participants in the µHEAT study.

**Table S2.** Per-sample metadata of stool samples in the µHEAT study.

**Table S3.** Exact p values and test statistics for all directed statistical tests reported in the manuscript.

**Table S4.** Somalogic custom panel of 1500 human serum proteins.

**Table S5.** Statistical results of MaAsLin2 linear models, serum proteins by ΔT category.

**Table S6.** Statistical results of MaAsLin2 linear models, serum metabolites by ΔT category.

**Table S7.** Statistical results of MaAsLin2 linear models, taxa and genes that differ temporally by ΔT category.

**Table S8.** Statistical results of MaAsLin2 linear models, transcriptional analysis by ΔT category.

**Table S9.** Statistical results of MaAsLin2 linear models, metagenomics analysis by ΔT category.

**Table S10.** Statistical results of MaAsLin2 linear models, metagenomics analysis by antibody category.

**Table S11.** GSEA test statistics for organoid RNA-seq.

**Table S12.** MEDI-detected foods that correlate with dietary category or Waltera abundance.

**Table S13.** Bacterial strains used experimentally in the manuscript.

